# *In vitro* anti cancer, anti inflammatory and anti oxidant activity of Celery oil (*Apium graveolens*)-Myristic acid based microemulsion system: Characterization and Biomedical application

**DOI:** 10.1101/2024.11.09.622815

**Authors:** Balaji Govindaswamy, Bharathi Ravikrishnan, Diptisree Panda, Sarath Perumal

**Author notes:** **Corresponding author: Balaji Govindaswamy,**. **Equal authorship:** Dipti Sree Panda and Sarath Perumal have contributed equally.

## Abstract

Plant based compounds are frequently modified, developed into drug components to treat different types of cancer. Celery is one of the most used ingredients in daily life, exhibits potential medicinal properties and myristic acid exhibits anti microbial and inflammatory properties making it ideal candidate for combinational therapy. This study demonstrates the formulation of celery oil-myristic acid based microemulsion system with sequence of weight/volume of components to optimize stable microemulsion. the sequence of formulations were performed, optimized through pseudo ternary phase diagram and stability analysis, of which CMA ME 4 formulation were observed to transparent and stable. This CMA ME 4 was analyzed using DLS for particle size which is found to be 14.0±27.1 and PDI 0.34±0.46 through 5th day to 30th day. In vitro drug release study has been performed with stimulated physiological condition (pH 5.6) and stimulated tumor tissue (pH 7.4), the microemulsion was withdrawn in time intervals and the kinetic model is constructed to detect the behaviour of the drug release profile. Further, anti inflammatory, anti diabetic and anti obesity assays were performed to analyse the potentiality of CMA ME 4 microemulsion formulation, where the results exhibited from 80 to 90% radical scavenging and inhibition compared with the synthetic drugs. A good cytotoxic activity of CMA ME 4 microemulsion was determined by positive results of the MTT assay performed with HeLa with 80 to 90% cell death, MCF7 with 80 to 90% cell death and HUVECs was used to determine the toxicity level when the microemulsion interacts with the normal cell which is found to be 80 to 90% cell viability over time. These findings more suitably suggest that combination of natural oil with synthetic components can be effective in treatment against diseases.

## Introduction

Modern medicine constant need for sophisticated drug delivery methods has prompted extensive research into natural substance, particularly fatty acids and essential oils, because of their inherent medicinal qualities, biocompatibility, and low risk of side effects. These natural agents, especially when used in nano-scale formulations, present a promising way to get around problems with traditional drug administration, such the restricted bioavailability, fast degradation, and low solubility of many therapeutic chemicals [1]. With a range of pharmacological actions, such as anti-inflammatory, anti-oxidant, and anti-bacterial properties, *Apium graveolens* (Celery) oil has become a powerful bioactive agent among natural oils [2]. Packed with volatile chemicals such as phthalides, β-selinene, and limonene, celery oil has been shown to have promise in treating infections, inflammatory disorders, and illnesses linked to oxidative stress [3]. Furthermore, adding essential fatty acids like myristic acid to formulations has practical advantages by improving the compounds lipophilicity and capacity to pass through biological membranes [4]. This creates a special chance to create a microemulsion based drug delivery system that combines myristic acid and celery oil [5]. Because of their transparent, isotropic, and thermodynamically stable properties, microemulsions have attracted a lot of attention in the field of nanomedicine [6]. Microemulsions, which are made up of water, oil and surfactants, have droplet sizes that are usually less than 100 nm [7]. This allows for improving drug solubilisation and greater bioavailability of medications that are not very water soluble. They are perfect carriers for hydrophobic medications because of their structural characteristics, which enable the integration of lipophilic substances like celery oil [8].

Additionally, because of their distinct interfacial characteristics and nanoscale droplet size, microemulsions offer a regulated release mechanism that enhances medication penetration and retention duration in the target tissues [9]. Microemulsions have the potential to be used in a number of biomedical domains, such as cancer treatment, dermatology, and the management of chronic inflammatory illnesses, since they are more able to pass through biological barriers than conventional formulations [10]. The formulation elements and circumstances must be carefully balanced when designing a microemulsion system for medication delivery [11]. To get the best possible balance between stability, transparency, and bioactivity, the oil, surfactant and co-surfactant ratios must be precisely adjusted [12]. Even while microemulsions are naturally stable, it can still be difficult to keep the system stable under different physiological circumstances, particularly when medication release rates need to be closely regulated to guarantee therapeutic effectiveness. Therefore, in order to successfully optimise the delivery of bioactive agents, a detailed knowledge of the interactions between these components is necessary for the creation of a celery oil-myristic acid microemulsion system [13]. Apart from their structural advantages, microemulsions provide regulated medication release according to environmental variables and pH sensitivity [14]. Applications in cancer, where tumor tissues frequently exhibit a slightly acidic environment, benefit greatly from this property.

To minimise systemic toxicity and reduce off target effects, the celery oil-myristic acid microemulsion system may be engineered to release medications selectively at particular places. By evaluating drug release in vitro under both pathological and physiological setting, we investigate this potential to see how these variables affect the microemulsion stability and drug release kinetics. We performed cytotoxic tests on HeLa and MCF7 cell lines to evaluate the medicinal potential and safety profile of celery oil-myristic acid microemulsion system. Celery oil is said to have anti-proliferative and pro-apoptotic qualities due to its rich content of bioactive chemicals including flavonoids and carotenoid [15]. These attributes helps explain its effectiveness against cancer cell lines by interfering with cellular signalling pathways, which are connected to growth and survival. Furthermore, myristic acid, which is known to improve membrane permeability and cellular absorption, may make it easier to transfer active ingredients straight to cancer cells [16], hence boosting the cytotoxic efficacy against resistance cell populations. The safety and selectivity of the microemulsion technology, we concurrently evaluated normal cytotoxicity utilising HUVECs [16]. In order to get insight into potential off-target effects on non-cancerous cells, HUVECs are frequently utilised to simulate the vascular endothelium. This work intends to create a therapeutic index for the celery oil-myristic acid microemulsion system by comparing the cytotoxic reactions of carcinogenic and non-cancerous cell types, demonstrating its suitability as a targeted and selective cancer treatment method. This provides a basis for creating medication delivery methods that are safe, efficient, and naturally generated while having influence on healthy cells. This work suggests creating and characterising a microemulsion system based on myristic acid and celery oil for use in biomedicine. This research help develop natural product based drug delivery drugs for therapies that need site-specific and regulated release.

## Methodology

### Materials

The Myristic acid and Celery oil were procured from Sigma Merck. The cell lines HeLa, MCF7 and HUVECs were purchased from NCCS Pune, India. The chemicals used in all the assays were procured from thermo scientific.

### Preparation and optimization of microemulsion

Conventional titration method was used for the preparation of microemulsion. Sequences of oil with surfactant/co-surfactant were mixed initially, titrated with double distilled water at 30±2°C, 7 variations were prepared and 1 with transparent appearance were chosen (CMA-ME 1-7) table 1. The formulations were formulated using myristic acid/ celery oil (0.10% w/w) as oil phase, sorbitan monolaurate as surfactant, ethanol as co-surfactant and double distilled water as aqueous phase. Ternary phase diagram software was used to construct the phase diagrams.

**Table 1:** Formulation sequence of microemulsions.

| Sequence code | Celery oil | Myristic acid % | Sorbitan monolaurate % | Ethanol % | Distilled water % |
| --- | --- | --- | --- | --- | --- |
| CMA-ME 1 | 0.10% w/w | 0.2 | 5 | 2 | 50 |
| CMA-ME 2 |  | 0.4 | 10 | 4 |  |
| CMA-ME 3 |  | 0.6 | 15 | 6 |  |
| CMA-ME 4 |  | 0.8 | 20 | 8 |  |
| CMA-ME 5 |  | 0.10 | 25 | 10 |  |
| CMA-ME 6 |  | 0.12 | 30 | 12 |  |
| CMA-ME 7 |  | 0.14 | 35 | 14 |  |

#### Characterization

##### Visual inspection & stability analysis

The CMA-ME 4 was initially observed for clear and stable appearance. The formulation was observed by eye using black backboards: clear, cloudy or sediment appearances were noted. The samples were eventually subjected to thermodynamic stability analysis, exposed to hot/heat temperature 40±50°C, room temperature 37±3°C, high stress test and cold temperature 33±40°C, ideally for 24, 48 and 72 hrs.

##### Particle size and PDI

Dynamic light scattering was utilized to measure the size and PDI (poly dispersity index). The CMA-ME 4 was initially diluted 10^5^ and subjected to measurement, triplicate readings were taken in account for evaluation (Malvern Zetasizer, UK) [17].

##### pH & Transmittance

The CMA-ME 4 subjected to pH meter. The pH rod were inserted and noted for fluctuations in determining the correct value. The transmittance of CMA-ME 1 to 7 was determined using a UV spectrophotometer at wavelength of 650 nm. Briefly 1ml of optimized CMA-ME 1 to 7 was added to the cuvette for determining the transmittance of light [18]. The sample analyzed in triplicate (n=3).

##### ATR-FITR

Active compound compatibility with the components of the emulsion was investigated by performing Attenuated Total Reflection-Fourier Transform Infrared Radiation (ATR-FTIR). ATR-FTIR analysis was performed on Celery-Myristic mix, S-mix and CMA-ME 4. A spectrum-two FTIR spectrometer (Agilent CARY 630) was used [19]. The transmittance range was from 500 to 4000 cm^-1^ with 4 cm^-1^ of resolution.

##### SEM

The morphological structure of optimized microemulsion was observed by scanning electron microscope operated using a Carl Zeiss EVO 18 at an accelerating voltage of approximately 5 kV. The average diameter was observed by Image J digital software analysis in different regions of the specimen [20].

##### Entrapment efficacy

The stabilized microemulsion was analyzed for drug loading efficiency; a 1 ml of CMA ME 4 was centrifuged at 15,000 RPM for 30 minutes at 25°C. The supernatant containing the un-encapsulated oil mix was screened using UV Spectrophotometer from 400 nm to 600 nm [21]. The absorbance reading was pointed against the calibration curve.

##### *In Vitro* Drug release and kinetics study

The in vitro release prolife of celery oil from the CMA ME 4 microemulsion was evaluated using a dialysis membrane in two different stimulant media: Physiological conditions (pH 7.4) and tumor tissues (pH 5.6). A 2 ml aliquot of the CMA ME 4 was placed into the dialysis membrane, sealed at both ends. The membrane was immersed in 50 ml of PBS (pH 7.4) and (pH 5.6), maintained at 37±0.5°C respectively. The medium was placed in orbital shaker at 100 rpm to maintain uniformity. At specific time intervals (0, 1, 2, 4, 6, 8, 12, and 24 hours), 2 ml samples were withdrawn from the release medium and replaced with an equal volume of fresh buffer to maintain uniform volumes. The concentration of the celery oil-myristic acid released into the medium was determined using UV Spec at 290 nm. The release data were fitted to kinetic models: zero order, first order, Higuchi and Korsmeyer Peppas for both the pH conditions to determine the release mechanism [22].

#### Biomedical application

##### DPPH assay

The DPPH radical scavenging activity was evaluated to ascertain the antioxidant capacity [23]. Various concentrations of the CMA ME 4, celery oil, myristic acid (10, 25, 50, 100, 200 µg/ml) were prepared. In brief, 1 ml of each concentration was mixed with 2 ml of freshly prepared 0.1 mM DPPH solution in methanol. The reaction mixture was incubated in the dark at RT for 30 minutes. The absorbance was measured at 517 nm using a UV Spectrophotometer (n=3). Ascorbic acid was used as a control, and the scavenging activity was calculated using the following equation:

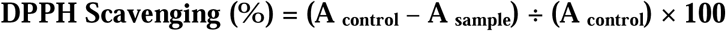

##### FRAP assay

The FRAP reagent was freshly prepared by mixing 300 mM acetate buffer (pH 3.6), 10 mM TPTZ (2,4,6-tripyridyl-s-triazine) in 40 mM HCl, and 20 mM FeCl_3_.6H_2_O in 10:1:1 ratio. A 100 µl aliquot of CMA ME 4, celery oil, myristic acid at concentrations (10, 25, 50, 100, 200 µg/ml) was mixed with 3 ml of FRAP reagent and incubated at 37°C for 30 mins (n=3). The absorbance of the resulting blue complex was measured at 593 nm. The results were expressed as µM Fe^2+^ equivalents per gram of sample.

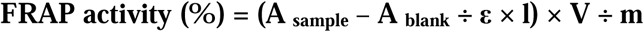

##### ABTS assay

The ABTS radical cation (ABTS+) was prepared [24] by mixing 7 mM ABTS solution with 2.45 mM potassium persulfate and allowing the mixture to stand in the dark at RT for 12-16 hours. Before use, the ABTS+ solution was diluted with ethanol to an absorbance of 0.70±0.02 at 734 nm. Series of concentrations CMA ME 4, celery oil, myristic acid (10, 25, 50, 100, 200 µg/ml) were added to 2 ml of ABTS+ solution and incubated for 30 mins at RT (n=3). Ascorbic acid was served as standard drug and the ABTS percentage of inhibition was expressed with the following formula:

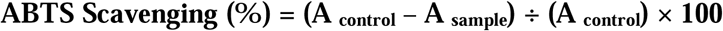

##### CUPRAC assay

To test the stability of antioxidant compounds, CUPRAC assay was performed [25]. The assay mixture was prepared by adding 1 mL of 10 mM copper (II) chloride, 1 mL of 7.5 mM neocuproine in ethanol, and 1 mL of 1 M ammonium acetate buffer (pH 7.0) in a test tube. To this mixture, 1 mL of the CMA ME 4, celery oil, myristic acid 4 (Conc. 10, 25, 50, 100, 200 µg/mL) was added. Trolox was used as standard reference with same concentrations. The final volume was adjusted to 4.1 mL with deionized water. The reaction mixture was incubated for 30 minutes at RT. Absorbance was measured at 450 nm using UV-Vis spectrophotometer (n=3).

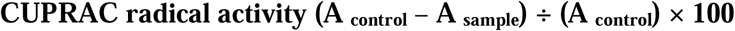

##### α-Amylase assay

The α-amylase inhibition activity of the ME [26] was assessed with CMA ME 4, celery oil and myristic acid (10, 25, 50, 100, 200 µg/ml) concentrations was prepared. A 500 µl aliquot of each concentration was mixed with 500 µl of 0.02 sodium phosphate buffer (pH 6.9) containing α-amylase enzyme (0.5 mg/ml). The mixture was pre-incubated at 37°C for 10 minutes. After pre-incubation, 500 µl of 1% starch solution in 0.02 M sodium phosphate buffer (pH 6.9) was added to each tube. The reaction mixture was incubated at 37°C for 10 minutes. The reaction was stopped by adding 1 ml of dinitrosalicylic acid (DNS) reagent, followed by heating in a boiling water bath for 5 minutes. After cooling, the mixture was diluted with 10 ml of double distilled water, and the absorbance was measured at 540 nm (n=3). Acarbose was used as the standard drug, and the percentage inhibition of the α-amylase activity was calculated using the following formula:

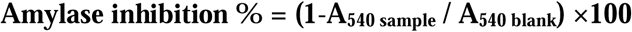

##### α-Glucosidase assay

The glucosidase inhibition assay was measured by monitoring the hydrolysis of p-nitrophenyl-α-D-glucopyranoside (pNPG), which releases p-nitrophenol [27]. The assay mixture consisted of 50 µl of α-glucosidase enzyme solution 0.1 U/mL in PBS (pH 6.8) and 50 µl of the CMA ME 4, celery oil, myristic acid and control at series of concentration (10, 25, 50, 100, 200 µg/mL). The mixture was pre-incubated at 37°C for 10 mins. After pre-incubation, 50 µl of 5 mM pNPG substrate solution was added to initiate the reaction. The procedure was carried out at 37°C for 30 mins and stopped by adding 50 µl of 0.1 M sodium carbonate. Acarbose was used as standard reference, measured at 420 nm using microplate reader (n=3). The following equation was used to calculate:

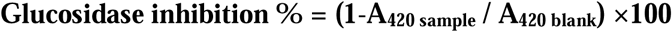

##### Protein denaturation inhibition assay

The anti-inflammatory activity of the ME was evaluated by assessing its ability to inhibit protein denaturation [28]. Various concentrations of the CMA ME 4, celery oil, myristic acid (10, 25, 50, 100, 200 µg/ml) were prepared for the assay, aspirin was used as control and diluted to same concentrations. A reaction mixture containing 1 ml of each concentration of ME, 1 mo of 1% (w/v) bovine serum albumin (BSA) solution, and 2 ml of PBS (pH 6.4) was prepared. The mixture was incubated at 37°C for 15 mins. Denaturation was induced by placing the reaction mixtures in a water bath at 70°C for 10 mins. After cooling to RT, the turbidity was measured at 660 nm using UV spectrophotometer (n=3). The percentage inhibition of protein denaturation was calculated using the following formula:

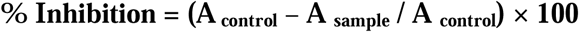

##### COX-2 enzyme inhibition assay

The COX-2 (Cycloxygenase) inhibition assay was performed to evaluate to anti-inflammatory [29]. The assay mixture was prepared by adding 150 µl of reaction buffer (Tris HCl, pH 8.0), 10 µl of COX-2 enzyme solution (0.1 U/mL), and 20 µl of CMA ME 4, celery oil, myristic acid with concentrations (10, 25, 50, 100, 200 µg/ml) into each well of a 96 well plate. Indomethacin, a known COX-2 inhibitor, was used as control at similar concentrations. The mixture was incubated at 37°C for 5 mins. After the pre-incubation, 10 µl of 1 mM arachidonic acid was added to initiate the reaction. The mixture was incubated again at 37°C for 30 mins to allow the reaction to proceed. The production of PGE_2_ was measured using a colorimetric method, where 200 µl of chromogenic reagent was added to stop the reaction and develop colour. After few minutes, the absorbance of the resulting solution was measured at 590 nm using a microplate reader (n=3) and was calculated using the below formula:

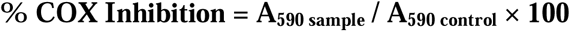

##### Lipase inhibition assay

The lipase inhibition assay was performed to evaluate the anti-obesity potential of the samples by measuring the inhibition of pancreatic lipase activity [30]. The assay mixture consisted of 200 µl of Tris-HCl buffer (pH 7.4), 50 µl of the pancreatic lipase enzyme solution (1 mg/ml), and 20 µl of the CMA ME 4, celery oil, myristic acid (10, 25, 50, 100, 200 µg/mL), Orlistat was used as control. The mixture was pre-incubated at 37°C for 15 mins. After pre-incubation, 20 µl of 5 mM pNPP solution was added to initiate the reaction. The reaction was carried out at 37°C for 30 mins. The hydrolysis of pNPP resulted in the formation of p-introphenol, and the absorbance of the reaction mixture was measured at 405 nm using a microplate reader and calculated using the below formula.

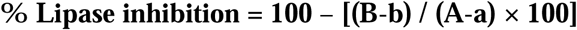

#### Evaluation of Cytotoxicity

##### Cell culture

HeLa (Human cervical adenocarcinoma) and MCF-7 (Human breast adenocarcinoma) was cultivated in DMEM (Gibco) supplemented with 10% FBS, 1.5 g/L Sodium bicarbonate and 1% penicillin-streptomycin (Gibco). Cells were maintained at 37°C in a humidified incubator with 5% CO_2_. HUVECs (Human Umbilical Vein Endothelial Cells) was cultured in DMEM (Gibco) supplemented with 10% FBS, 1% Penicillin-Streptomycin, and 1% Glutamine. The cells were maintained in humidified incubator with 5% CO_2_. Before experiments, cultures reached approximately 80% confluence, cells were harvested using trypsin-EDTA 0.25% (Gibco) and plated at an initial density of 5×10^3^ cells/well in 96 well plate for further assays,

##### MTT assay and cell viability

The MTT assay was employed to evaluate the in vitro cytotoxic effects of CMA-ME 4, Celery oil, and Myristic acid on HeLa, MCF-7, and HUVECs cell lines. 5-Fluorouracil was used as the standard reference. Initially, 5×10^4^ cells per well were carefully placed in 96 well plate. These cells were exposed to varying concentrations of the CMA-ME 4, Celery oil, and Myristic acid (20-100 µg/mL). The cells were incubated for 24 hrs, after incubation, the culture media was replaced with 100 µl per well of a PBS solution containing MTT at a concentration of 0.5 mg/mL. The absorbance was measured at 490 nm using microplate reader (n=3).

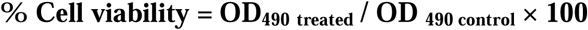

### Statistical analysis

The results were presented by performing statistical analysis using graph pad prism version 10.1.2.

## Results and discussion

### Ternary phase and visual appearance

The ternary phase diagram predicts out the concentration range of components to stabilize microemulsion easily. The Figure 1A show the pseudo-ternary of 7 sequence formulations each having different weight respectively, the shaded areas represents the clear/transparent and the stable emulsion was observed in CMA ME 4 after preparation. The Figure 1B shows the microemulsion blend of CMA ME 4 which is thermodynamically transparent; therefore, this microemulsion was optimized for further studies. The optimized microemulsion was observed with Sudan black dye staining for observing the lipid layers formed. The relationship among three elements: oil mix, Surfactant mix and water, the development of a stable, uniform phase, and homogenous. When hydrophilic and lipophilic components are properly balanced, microemulsions usually develops in ternary systems is stabilised by the S mix, which lowers the interfacial tensions between the water and the Oil mix. These immiscible phases can coexist in a stable emulsion because the surfactant molecules most likely align themselves at the oil-water contact. Myristic acid, a saturated fatty acid, adds to the lipophilic character of this phase, whereas other hydrophobic components make up celery oil. They help create a thermodynamically stable microemulsion when combined with water and surfactants [31].

**Figure 1:**
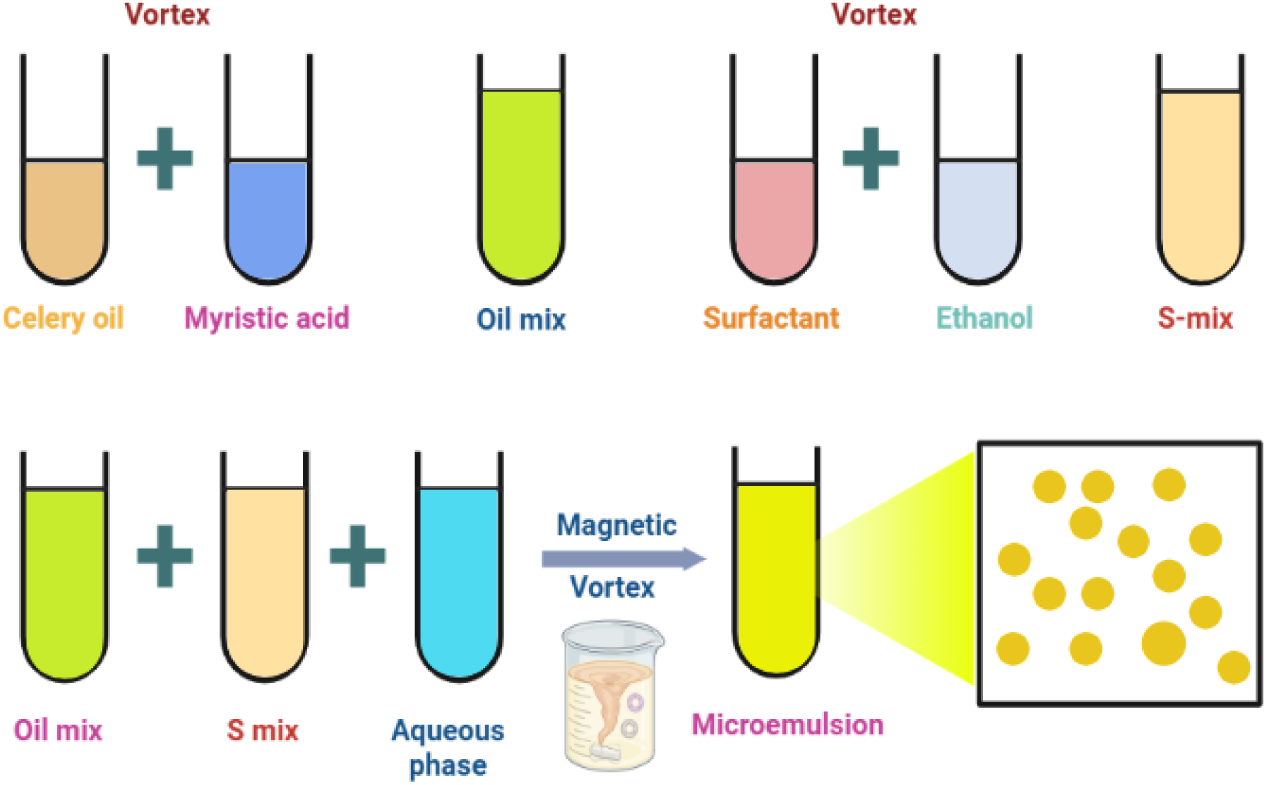
Schematic illustration of microemulsion synthesis

**Figure 2:**
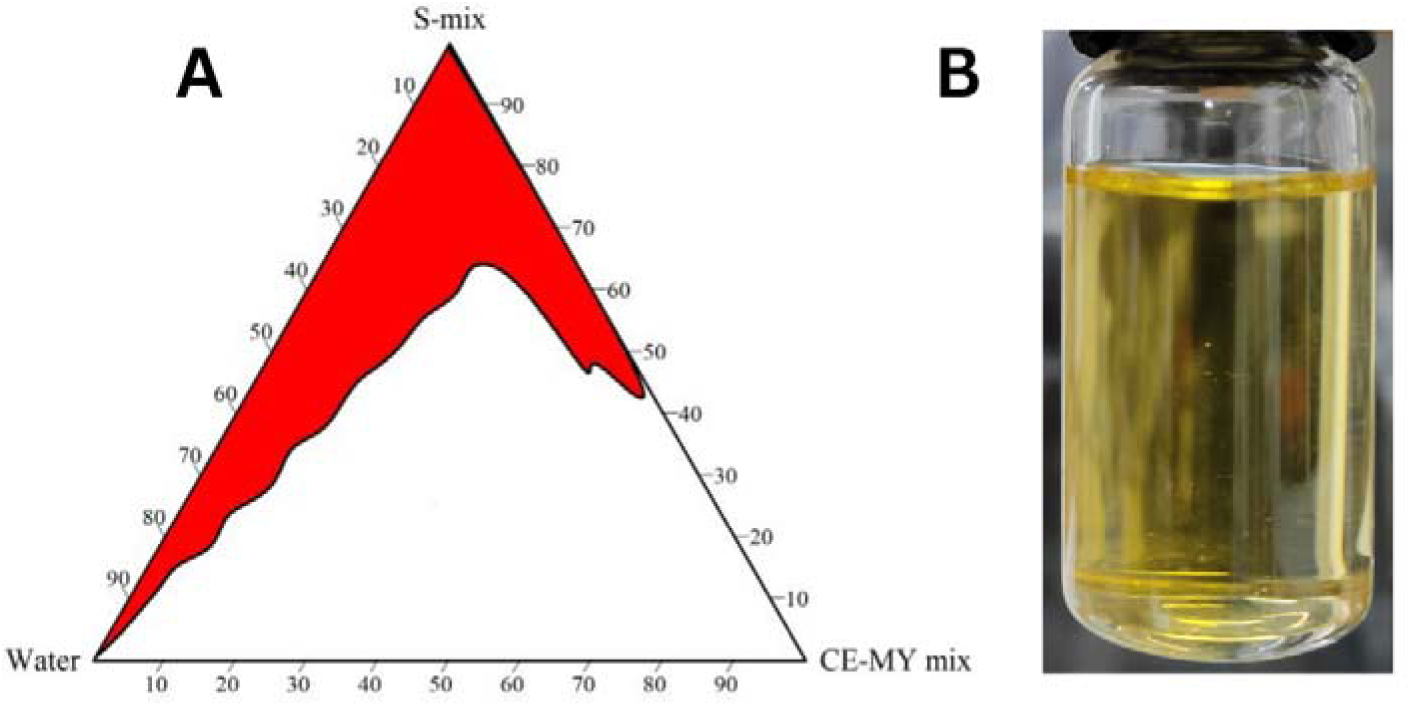
(A) Pseudo-ternary phases diagram of CMA ME 1 to CMA ME 7 (B) Visual appearance of CMA ME 4 (optimized).

### Stability analysis and DLS

The series of microemulsions post prepared are subjected to thermodynamic stability analysis, where these emulsions are exposed to heating, cooling and force stress studies to visually interpret that these microemulsions withstand eventually in any shelf-life condition. Table 2 depicts the thermodynamic studies implied on (CMA ME 1 to CMA ME 7) for 24, 48 and 72 hours. The visual inspection noted after the preparation for all these microemulsions were stable at 24 hours, during the completion of 48 hours CMA ME 3 and 4 showed stable microemulsion when compared with other microemulsions where the oil mix and surfactant mix formed a distinct thick mat layer. The CMA ME 5 to 6 was highly viscous and unstable. The heating test showed good results to the emulsions which are prepared with minimum amount of surfactant mix. The CMA ME 4 effectively showed good stability over time, which later was observed same in cooling test. Further, the microemulsion sequences were applied to freezing, thawing and high force stress test, CMA ME 4 showed better results when compared to other microemulsions. The DLS results for CMA ME 4 provide insights into both the particle size evolution and the polydispersity index (PDI) over a period of 30 days [32]. In Figure 3 (A) and (B) day 5, the particle size is 14.0 nm, indicating the initial formation of relatively small droplets within the microemulsion, and the PDI is 0.341, indicating a relatively narrow size distribution, which is a characteristic of a stable system with uniform particles. Day 10 shows the 23.6 nm particle size and PDI 0.452, which suggests growth or swelling of droplets over time, likely due to increased interactions or changes in the microenvironment of emulsion. Day 15-20, the particle size stabilizes around 25 nm, indicating that the system reaches a steady state with minimal further growth, PDI decreases slightly 0.394 and 0.396, reflecting a uniform size distribution compared to Day 10. Day 25-30, a slight increase in observed 26.1 to 27.1 nm showing that the system remains mostly stable but undergoes a small degree of further droplet growth. The PDI drops further to 0.328, indication improved homogeneity within the system. This suggests that the microemulsion becomes more uniform as it stabilizes over time. On day 30^th^, the PDI increases to 0.465, possibly indicating some degree of instability or the presence of larger aggregates within the system towards the end of the observation period [33].

**Figure 3:**
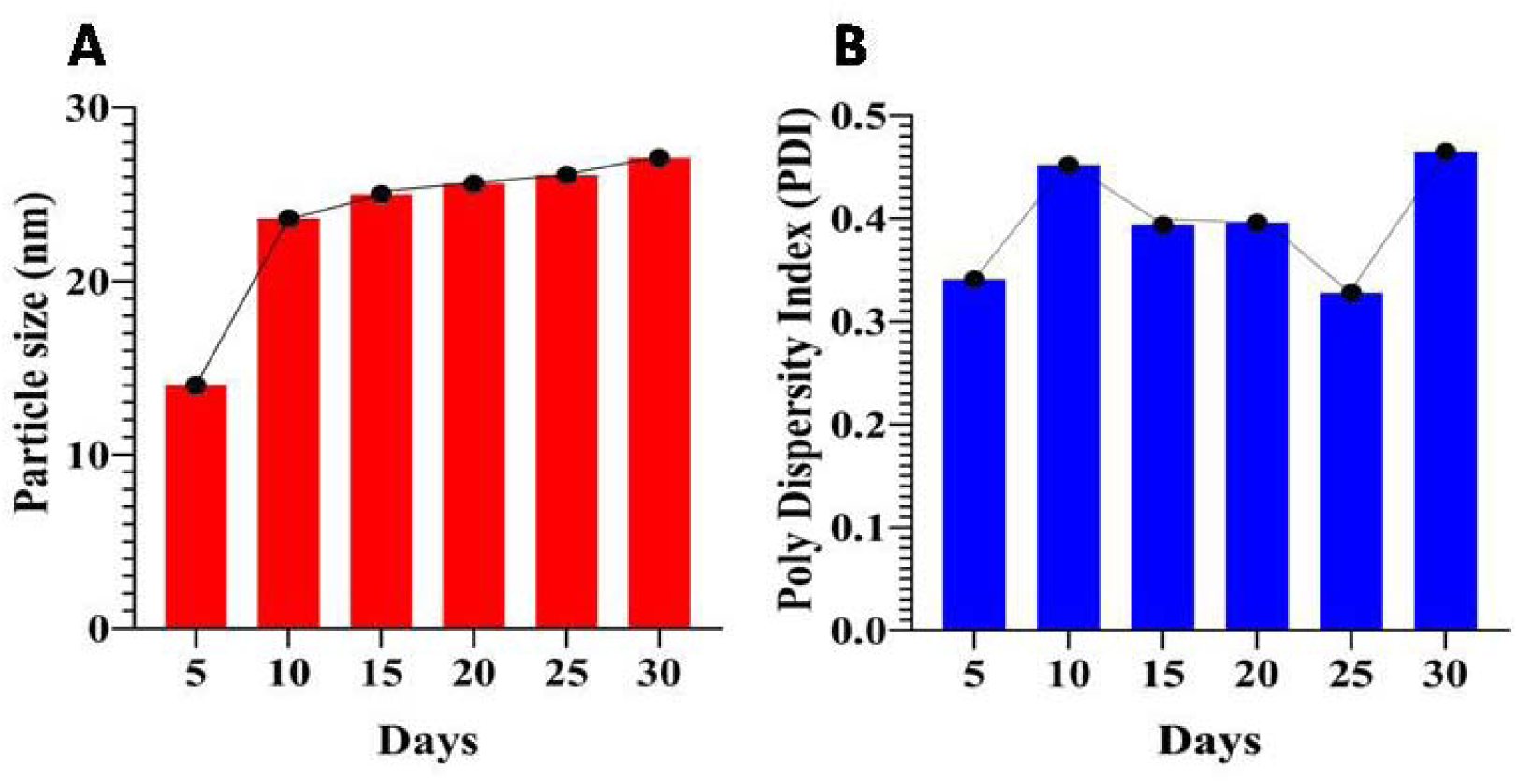
(A). Particle size analysis of CMA ME 4 (B). Poly Dispersity Index of CMA ME 4.The DLS assay performed over a month with 5 days increasing.

**Table 2:**
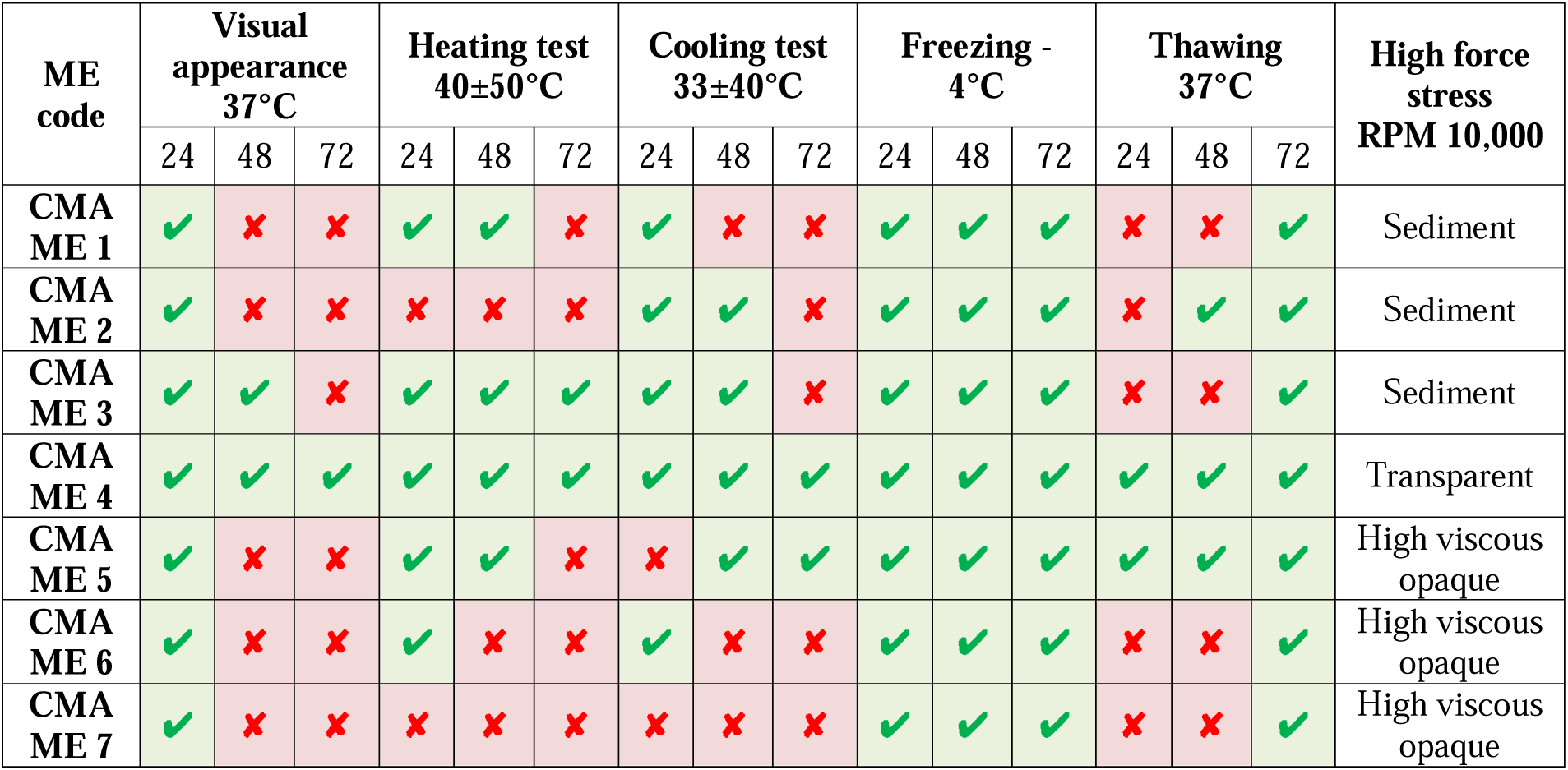
Stability analysis of CMA ME 1 to CMA ME 7 microemulsion sequence.

| ME code | Visual appearance 37°C |  |  | Heating test 40±50°C |  |  | Cooling test 33±40°C |  |  | Freezing - 4°C |  |  | Thawing 37°C |  |  | High force stress RPM 10,000 |
| --- | --- | --- | --- | --- | --- | --- | --- | --- | --- | --- | --- | --- | --- | --- | --- | --- |
|  | 24 | 48 | 72 | 24 | 48 | 72 | 24 | 48 | 72 | 24 | 48 | 72 | 24 | 48 | 72 |  |
| CMA ME 1 | ✓ | ✗ | ✗ | ✓ | ✓ | ✗ | ✓ | ✗ | ✗ | ✓ | ✓ | ✓ | ✗ | ✗ | ✓ | Sediment |
| CMA ME 2 | ✓ | ✗ | ✗ | ✗ | ✗ | ✗ | ✓ | ✓ | ✗ | ✓ | ✓ | ✓ | ✗ | ✓ | ✓ | Sediment |
| CMA ME 3 | ✓ | ✓ | ✗ | ✓ | ✓ | ✓ | ✓ | ✓ | ✗ | ✓ | ✓ | ✓ | ✗ | ✗ | ✓ | Sediment |
| CMA ME 4 | ✓ | ✓ | ✓ | ✓ | ✓ | ✓ | ✓ | ✓ | ✓ | ✓ | ✓ | ✓ | ✓ | ✓ | ✓ | Transparent |
| CMA ME 5 | ✓ | ✗ | ✗ | ✓ | ✓ | ✗ | ✗ | ✓ | ✓ | ✓ | ✓ | ✓ | ✓ | ✓ | ✓ | High viscous opaque |
| CMA ME 6 | ✓ | ✗ | ✗ | ✓ | ✗ | ✗ | ✓ | ✗ | ✗ | ✓ | ✓ | ✓ | ✗ | ✗ | ✓ | High viscous opaque |
| CMA ME 7 | ✓ | ✗ | ✗ | ✗ | ✗ | ✗ | ✗ | ✗ | ✗ | ✓ | ✓ | ✓ | ✗ | ✗ | ✓ | High viscous opaque |

### Physical characterizations and ATR-FTIR

The optimized microemulsion was subjected to physical characterization like as pH, and transmittance. The pH profile Figure 4 (A), of the CMA ME 4 microemulsion demonstrates a time-dependent acidification over the course of 25 days. initially, on day 5, the pH is recorded at approximately 5.7, which steadily decreases to about 5.4 by day 25. this decline in pH could indicate hydrolytic degradation of the emulsion components, which often occurs in microemulsion systems over time. this acidification trend might affect the stability, efficacy and biocompatibility of the formulation [34]. The transmittance profile of the microemulsion sequence Figure 4 (B) exhibited a bell shaped curve, with the CMA ME 4 demonstrating the high transmittance (close to 100%), signifying a transparent formulation. This peak transparency indicates a well-dispersed, stable microemulsion with nanosized droplets, ensuring minimal light scattering. In contrast, CMA ME 1-3, and 7 showed significantly lower transmittance, reflecting cloudy or opaque nature. This clarity is attributed to phase separation, larger droplet size, or incomplete emulsification in these formulations [35]. FTIR analysis Figure 4 (C) was done and the results obtained were in accordance with reported literatures. Each peak indicates the absorption that occurred as a result of the presence of distinct functional groups in each of the samples. A strong, sharp peak near 2853 cm^−1^indicates the presence of symmetric CH_2_stretching. Another peak near 2922cm^−1^ suggests asymmetric CH_2_ stretching. The presence of CH3 groups might be indicated by a peak around 2872 cm^−1^ (symmetric) and 2959 cm^−1^ (asymmetric).The presence of aromatic rings in the oil mix can be indicated by peaks in the 1600-1450 cm^−1^ region, corresponding to C=C aromatic ring stretching. Depending on the specific oil type, functional groups like esters (C=O stretching around 1740 cm^−1^) or ketones (C=O stretching around 1715 cm^−1^) might be present and show up in the spectrum. The S-mix displayed a slight peak around 3473 cm^−1^ signifying O–H stretching and defined peaks at 2876 cm^−1^ and 2857 cm^−1^ representing methylene stretching vibrations. It also displayed a sharp peak at 1092 cm^−1^ which shows the stretching vibration of –CH_2_–O–CH_2_– and a peak at 1737 cm^−1^ signifying the presence of carbonyl group from R–CO–O–R. The water phase in the microemulsion caused a long stretching peak from 3000 cm^−1^ to 3500 cm^−1^ representing the presence of OH group [36,37].

**Figure 4:**
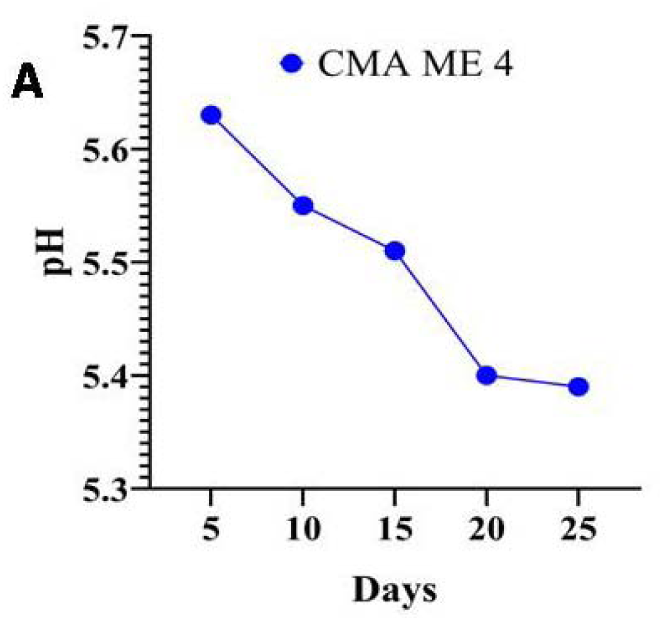
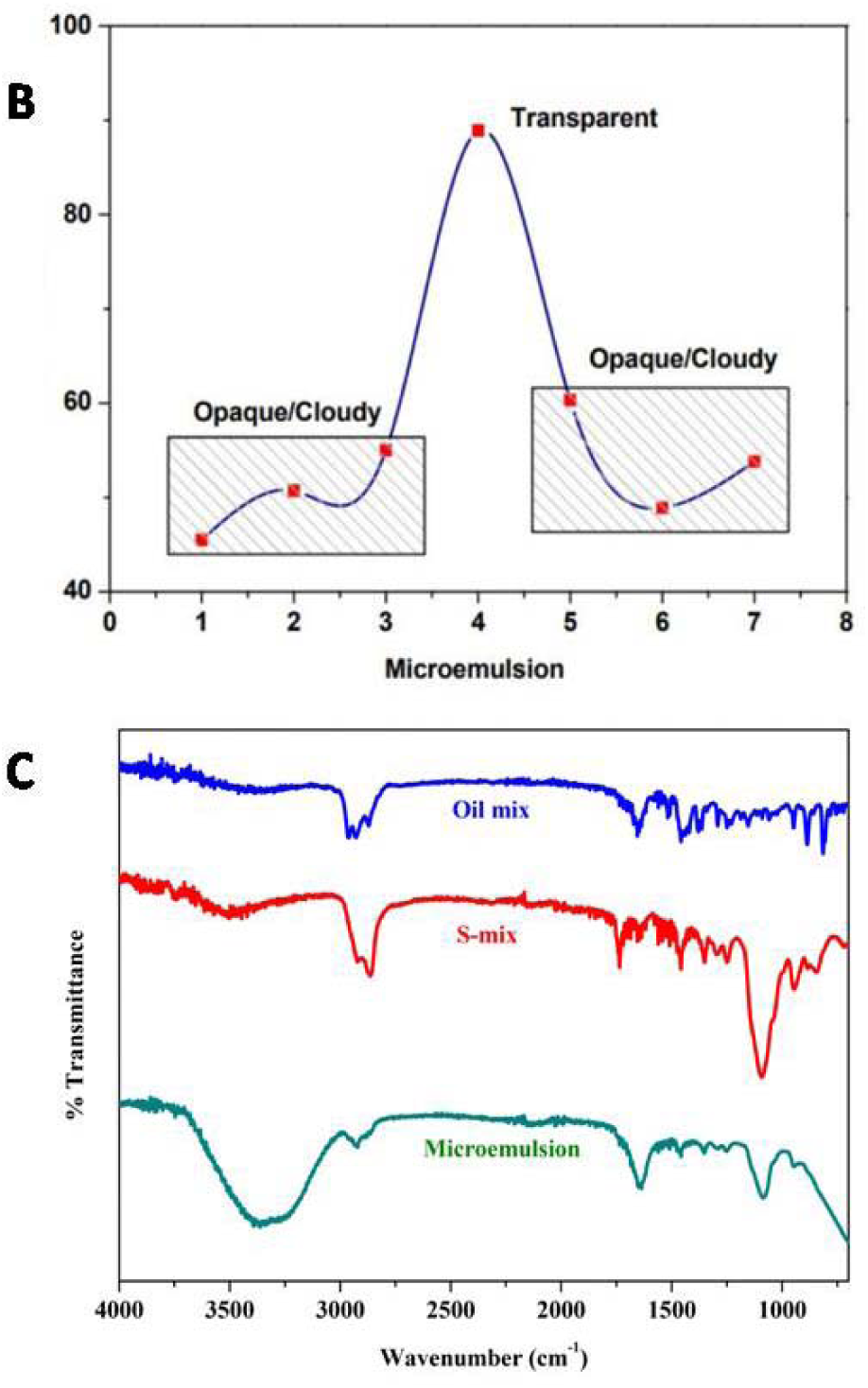
(A) pH analysis (B) Transmittance using UV Spec (C) ATR-FTIR analysis of optimized microemulsion, S mix and O mix

### SEM analysis

SEM images were obtained using a Carl Zeiss EVO 18 model SEM, providing high-resolution images of the emulsion. Across images, notably in Figure 5 (A) and Figure 5 (D), we observe a pattern of uniformly dispersed, spherical droplets embedded within the matrix. This distribution indicates the effective incorporation of celery oil, which, rich in essential fatty acids and antioxidants, contributes to the therapeutic efficacy of the formulation. The uniform droplet size observed in these images suggests that the celery oil is evenly distributed throughout the emulsion, reducing the risk of aggregation or phase separation. SEM images Figure 5 (E) and Figure 5 (G) reveal a smooth, cohesive surface morphology with minimal droplet coalescence. The consistent droplet size observed in these images highlights the stabilizing effect of myristic acid, which prevents aggregation and supports the structural integrity of the microemulsion. This characteristic is essential for preserving emulsion stability over time, as it reduces the likelihood of phase separation. The fine dispersion pattern in Figure 5 (B) and SEM-Liq-A-08 underscores the formulation’s high surface area, crucial for maximizing the bioavailability of hydrophobic compounds in celery oil. The images show that the emulsion achieves optimal dispersion without compromising structural integrity, which could facilitate improved delivery and interaction with biological membranes. This is particularly valuable in therapeutic applications, where efficient delivery of active compounds is paramount. The uniform globule morphology observed across images (notably Figure 5 (A), Figure 5 (D), Figure 5 (C), and Figure 5 (F) highlights a stable emulsion structure, while the high surface area observed in Figure 5 (B) and Figure 5 (H) suggests potential for enhanced bioavailability [38,39].

**Figure 5:**
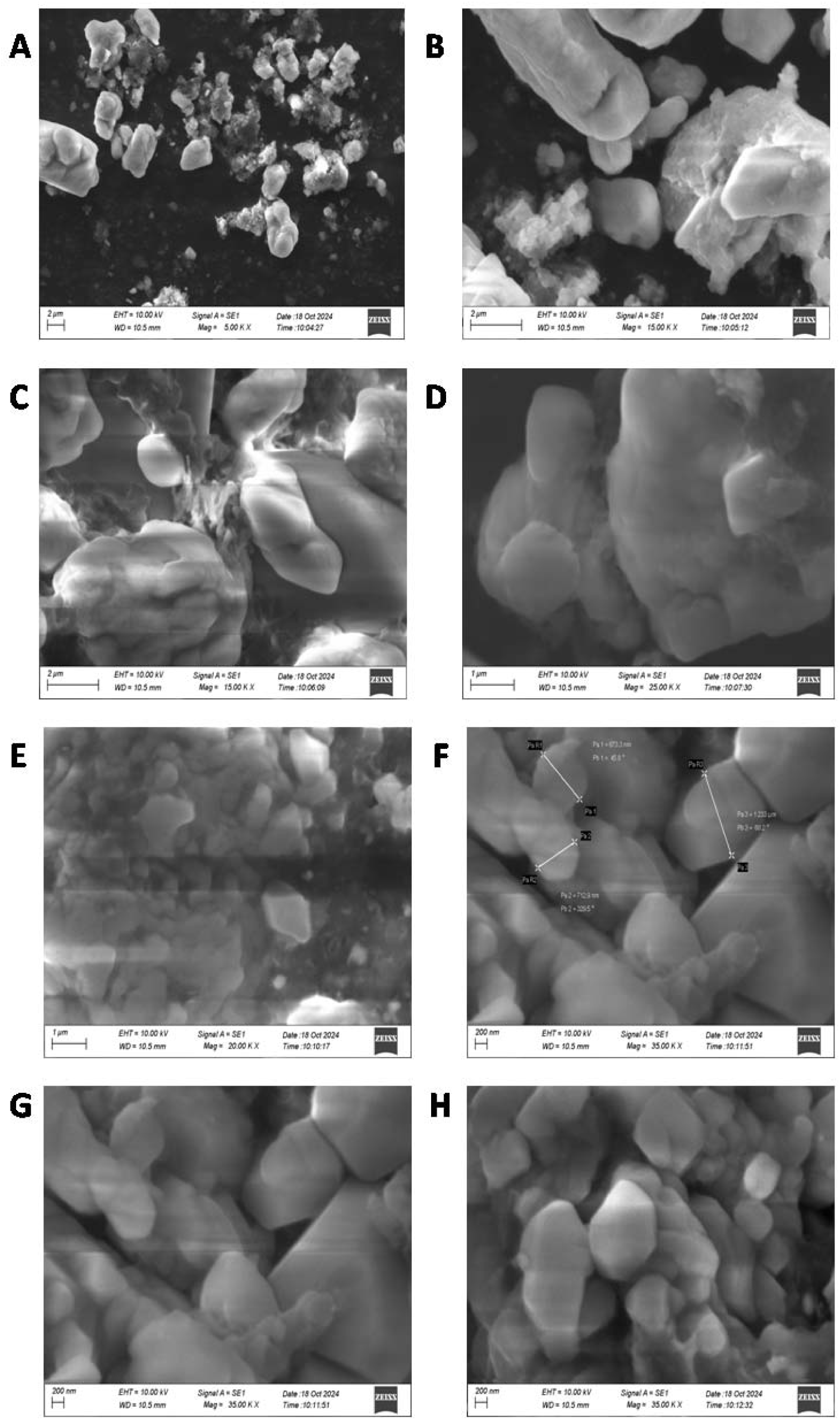
SEM analysis of optimized microemulsion CMA ME 4

### Entrapment efficiency and *In vitro* drug release with kinetic study

The entrapment efficiency data Figure 6 (A) presents a strong linear relationship between absorbance and drug concentration, with an R^2^ value of 0.9971, indicating precise quantification of the oil mix within the microemulsion. The *in vitro* drug release profile in both pH 7.4 (stimulated physiological condition) and pH 5.6 (stimulated tumor tissue conditions) demonstrate sustained release over a 24 hour period, with notable differences in the kinetics observed under each condition. In Figure 6 (B) at pH 7.4, the release follows a zero-order kinetic model, suggesting that the drug release is independent of concentration and occurs at a constant rate, which is desirable for achieving a stable therapeutic effect. The first-order model, which implies concentration dependent release, also fits the data, indicating that some portion of the release may be influenced by the drug concentration gradient. The Higuchi model, which describes release through diffusion from a matrix, and the Korsmeyer-Peppas model, which characterizes anomalous (non-fickian diffusion, are both applicable, particularly during later stages of the release profile, reflecting a combination of diffusion-controlled mechanisms. In contrast, the release profiles, at pH 5.6 Figureure 6 (B) shows a slower and more controlled release. The Korsmeyer-Peppas and Higuchi models describes the release kinetics, suggesting that the release mechanism is primarily diffusion-driven under these conditions, possibly due to pH dependent changes in the microemulsion system or drug solubility. The zero order less favourable fit indicates that the release in this environment is not constant but rather influenced by diffusion and potential structural alterations of the microemulsion at lower pH. Myristic acid and celery oil may work in concert to stabilise the microemulsion and guarantee that it is compatible with biological membranes. Bioactive substances found in celery oil may improve the solubility and bioavailability of hydrophobic medications, and myristic acid may help the systems lipid compatibility, which would assist drug release even more. This combination probably improves the emulsions physical stability and drug delivery capabilities. CMA ME 4 is a viable option for pharmaceutical applications due to its stability and prolonged release behaviour, particularly in situations for controlled release, such the treatment of cancer or chronic diseases. Strong physicochemical stability, which is essential for practical therapeutic applications, is suggested by the stability offered by the optimised formulation, particularly given that CMA ME 4 is extremely clear [40,41].

**Figure 6:**
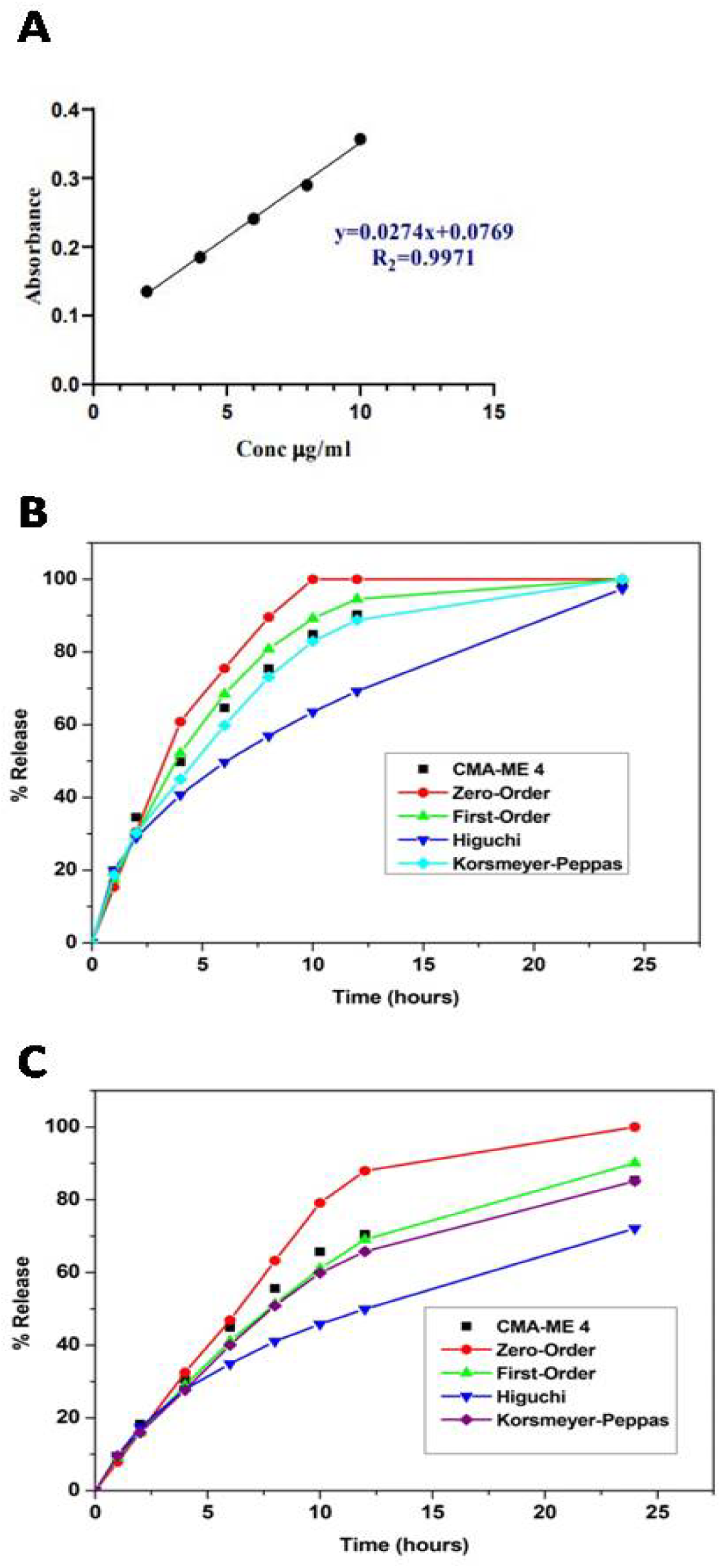
(A) Entrapment efficiency (B) in vitro drug release profile in pH 7.4 (C) In vitro drug release profile in pH 5.6

### Antioxidant assay

The DPPH assay was employed to evaluate the free radical scavenging ability of the celery oil-myristic acid microemulsion CMA ME 4 compared to its individual components celery oil, myristic acid and a standard antioxidant, ascorbic acid. In Figure 7 (A), the radical scavening acitivty for all the samples increased with concentration ranging from 10 to 200 µg/mL. At the highest concentration 200 µg/mL, CMA ME 4 demonstrated superior radical scavenging activity 78.3% compared to celery oil 66.7% and myristic acid. Figure 7 (B) illustrates the activity of FRAP at the 5^th^ concentration, CMA ME 4 exhibited highest FRAP value, outperforming both celery oil and myristic acid, trolox, a reference antioxidant displayed the highest reducing activity similarly matching with CMA ME 4. The enhanced DPPH and FRAP activity of CMA ME 4 suggests a significant increase in antioxidant capacity due to combined effects of celery oil and myristic acid. The ABTS^+^ radical cation assay Figureure 7 (C), demonstrates the highest concentration exhibits the highest scavenging activity CMA ME 4 89.5%, celery oil 74.8%, myristic acid 70.2% and ascorbic acid 93.2%, the CMA ME 4 has the potential activity, due to improved solubility and dispersion of components in microemulsion system. The CUPRAC assay Figureure 7 (D) was employed to asses the cupric ions Cu^+^ reducing antioxidant capacity of the samples, at highest concentrations CMA ME 4 exhibited 78.8%, celery oil 65.3%, myristic acid 61.8% and trolox showed highest value 82.4%. Multiple free radical scavening and reducing power studies show that CMA ME 4 components such as celery oil and myristic acid has strong antioxidant activity, as indicated by the results of all assays. The formulation makes use of the inherent antioxidant qualities and the microemulsion amplifies these benefits through enhanced solubility, stability, and perhaps synergistic interactions. Because of this, CMA ME 4 is a good option for potent antioxidants, such pharmaceutical and nutraceutical compositions meant to treat illnesses linked to oxidative stress [42,43].

**Figure 7:**
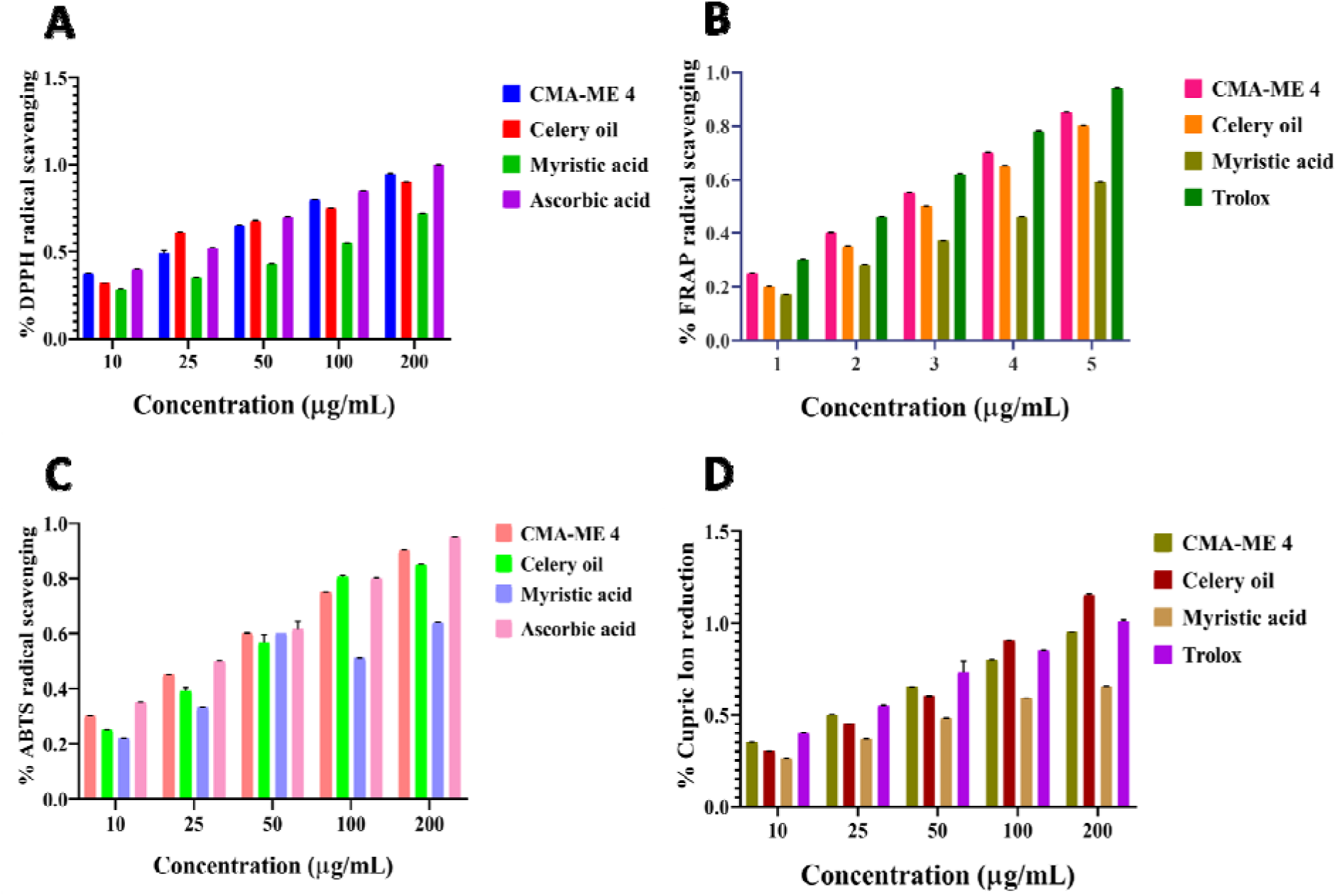
Anti oxidant assays (A) DPPH (B) FRAP (C) ABTS (D) CUPRAC of CMA ME 4, celery oil and Myristic acid

### Antidiabetic activity

In the α-amylase inhibition assay Figure. 8 (A), CMA ME 4 reaches a peak inhibition of approx 75% at 200. In contrast, celery oil and myristic acid shows increasing and decreasing inhibition levels. In Figure 8 (B), α-glucosidase assay CMA ME 4 exhibited good inhibition 80% at the highest concentration. Celery oil and myristic acid showed lower inhibition peaks ranging from 50-60%, the CMA ME 4 effectively worked in α-glucosidase assay, when compared with Acarbose, and it showed good inhibition 0.9 in both the assay showing good peaks. The combined findings of the α-amylase and α-glucosidase inhibition tests indicate that CMA ME 4 is useful in lowering the absorption of glucose and the digestion of carbohydrates. The stabilised, bio available ingredients in the CMA ME 4 may improve the formulations effectiveness by enabling reliable enzyme engagement. This makes CMA ME 4 a viable option for creating anti diabetic natural microemulsion treatment [44].

**Figure 8:**
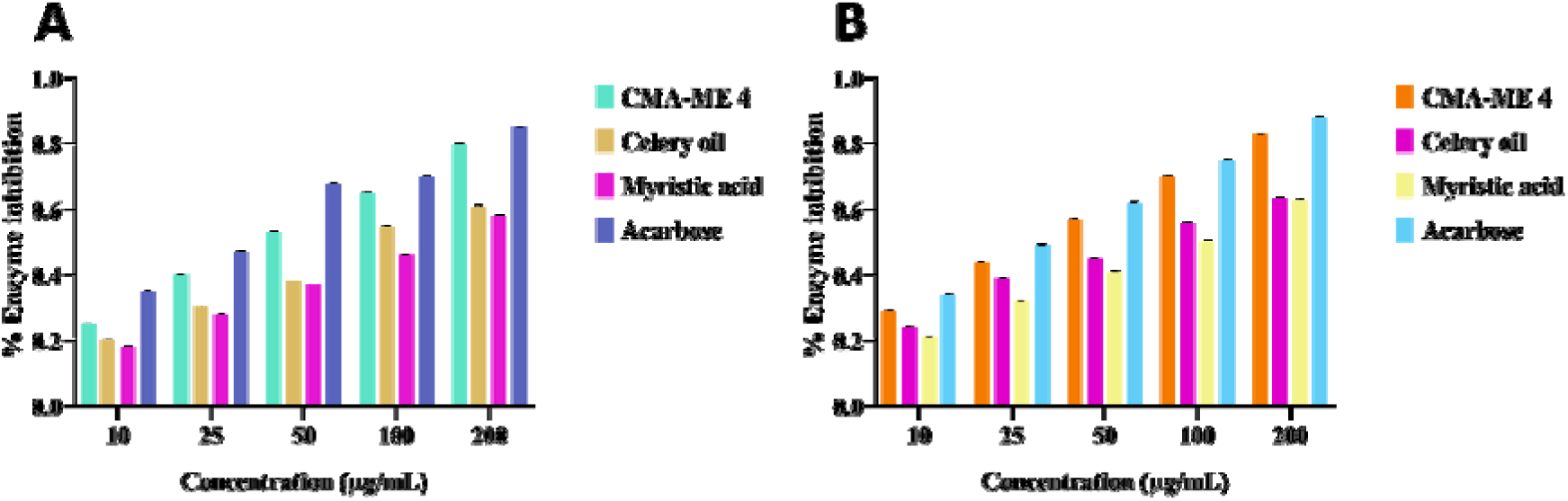
Anti-diabetic activity (A) α-amylase assay (B) α-glucosidase assay of CMA ME 4, Celery oil, Myristic acid and Acarbose.

**Figure 9:**
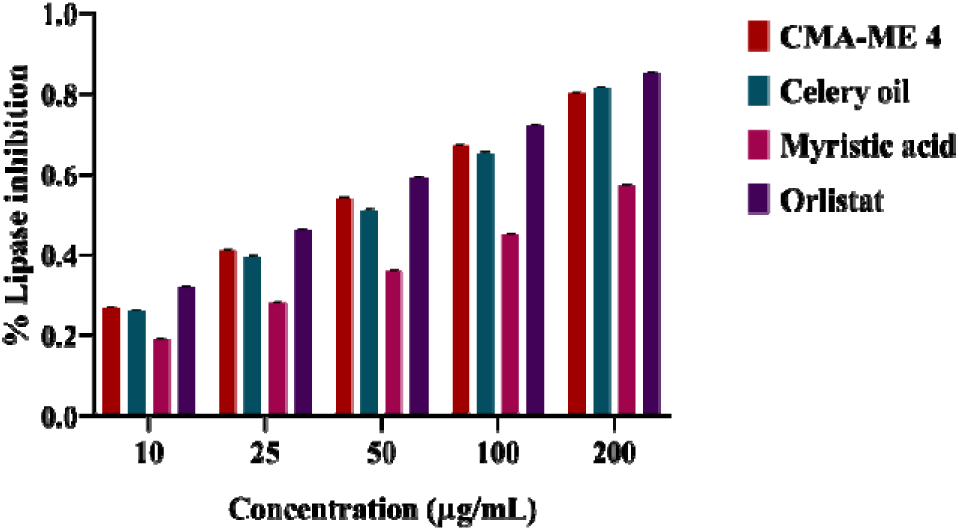
Anti-obesity assay of CMA ME 4, celery oil, myristic acid and Orlistat

### Anti-obesity

The lipase inhibition assay results present inhibitory effects of CMA ME 4, celery oil, myristic acid and Orlistat on pancreatic lipase activity. The data Figureure 9 reveals a concentration dependent increase in lipase inhibition across all tested sample, Orlistat showing a highest concentration as a standard, whereas CMA ME 4 and Celery oil shows equal inhibition peaks, which suggests that the oil and emulsion contain flavonoids and other bioactive essentials which works effectively in lipase inhibition. The myristic acid show lower inhibition peaks compared to others. Although myristic acid and celery oil each have qualities that can help inhibit lipase on their own, their combination in a microemulsion system may enhance these benefits. The fatty acid structure of myristic acid may have a direct impact on enzyme function, whereas bioactive chemicals found in celery oil interact with lipid metabolism pathways. These ingredients work better together in the microemulsion because of their increased stability, which allows for more reliable and consistent lipase inhibition [45].

### Anti-inflammatory

The protein denaturation assay and COX inhibition assay evaluates the potential of the celery oil-myristic acid test compounds to inhibit inflammation. Figure 10 (A), is the inhibition percentages increase as the concentration rises from 10 µg/ mL to 200 µg/ mL, Aspirin shows the highest inhibition at all concentrations, with around 90% inhibition at 200 µg/ mL. Celery oil and myristic acid shows moderate inhibition at higher concentration with value of 80%, whereas CMA ME 4 shows lower inhibition compared to the individual components, starting relatively lower at lower concentrations but rising steadily in higher concentrations 60%. In Figure 10 (B), COX inhibition assay, indomethacin shows 100% inhibition rate, similarly the celery oil reached 80-90% inhibition rate and myristic acid show moderate inhibition levels ranging from 50-70% at 200 µg/ mL. Both the anti inflammatory assays demonstrate that celery oil has a strong anti-inflammatory effect, which is further complement by myristic acid. The microemulsion CMA ME 4, while not exceeding the individual effects of its components, shows consistent inhibition in both assays. This could suggest that the microemulsion offers a sustained release or enhanced bioavailability that could be effective over time, despite slightly lower peaks. The results of COX inhibition and protein denaturation suggest that CMA ME 4 may function as a microemulsion based, natural anti inflammatory drugs. Because of its improved stability and synergistic effects, the microemulsion can be used in substitute therapies [46].

**Figure 10:**
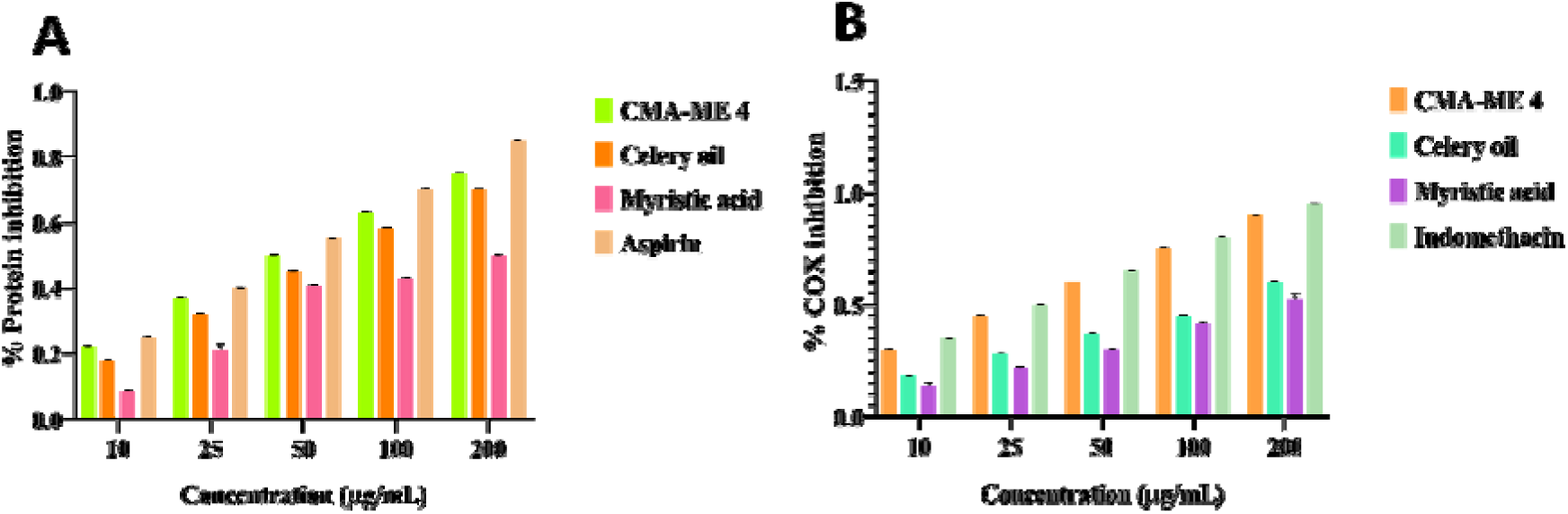
Anti-inflammatory assay (A) Protein denaturation assay (B) COX-2 inhibition assay

### MTT assay

The cytotoxicity was performed with three different cell lines, HeLa (cervical cancer), and MCF7 (breast cancer). Figure 11 (A), shows the cytotoxicity effects on HeLa and Figure 11 (B) displays the results for MCF7 cells. In HeLa cells, CMA ME 4 demonstrated significant cytotoxicity at concentrations above 40 µg/ mL, reducing cell viability to approx 50% and further decreasing to below 30% at 100 µg/ mL, as in MCF7 cells, a similar dose-dependent reduction in viability was observed, with around 60% viability at 40 µg/ mL and less than 25% at 100 µg/ mL. Celery oil displayed moderate anticancer activity in both cell lines. At 40 µg/ mL, viability was reduced to around 70% in HeLa cells, and 60% in MCF7 cells. At 100 µg/ mL, viability was further reduced to approximately 40% in HeLa and 35% in MCF7 cells demonstrating a lower potency compared to CMA ME4. Myristic acid had a mild cytotoxic effect on both the cell lines, at 40 µg/ mL, viability remained above 70% in HeLa cells and 75% in MCF7 cells. At the highest concentration 100 µg/ mL, viability was reduced to approximately 50% in HeLa cells and 55% in MCF7 cells, indicating that myristic acid alone is less effective than the microemulsion or celery oil. As expected, 5-fluorouracil exhibited potent anticancer activity, reducing cell viability to approximately 25% at 40 µg/ mL in both HeLa and MCF7 cells. At 100 µg/ mL, viability was reduced to below 20% confirming the efficiency of this chemotherapeutic agent as standard. The results indicate that the CMA ME 4 demonstrates enhanced anticancer activity compared to its individual components in both

**Figure 11:**
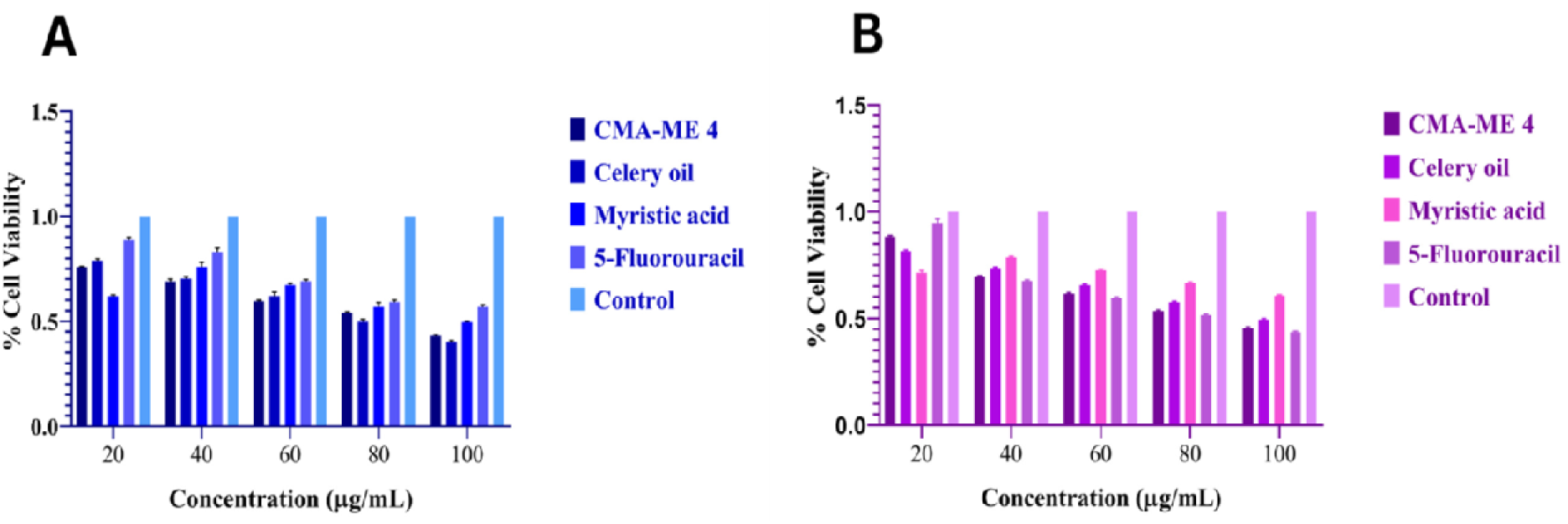
Anti-cancer MTT assay (A) HeLa cell line activity (B) MCF7 cell line activity

HeLa and MCF7 cells. CMA ME 4 ability to reduce cell viability to 30% at higher concentrations suggests its potential as a more effective anticancer treatment than its individual components. These findings suggest a synergistic effect between celery oil and myristic acid within the microemulsion system [47,48].

The cytotoxicity of CMA ME 4 shows the control group consistently maintained high cell viability across all concentrations tested, the CMA ME 4 group showed slightly decreased cell viability compared to the control group, but the reduction remained minor and consistent across all concentrations. In Figure 12, suggests that CMA ME 4 exhibits a lower level of cytotoxicity on HUVECs, as the cell viability remains above 80% even at the highest concentration 10. Similar to CMA ME 4, celery oil demonstrated only a minimal reduction in cell viability relative to the control group, including low toxicity. However, there appears to be a slight decrease in cell viability with increasing concentration. Cell viability consistently remains above 80%, suggesting that celery oil, on its own, does not exhibit significant cytotoxicity. Myristic acid showed a more reduction in cell viability compared to CMA ME 4 and celery oil, especially at higher concentrations. Despite this cell viability remained generally above 75% across all concentrations, indicating low to moderate cytotoxicity in HUVECs. This low level cytotoxicity across all test groups suggests that CMA ME 4 and its components are biocompatible with HUVECs, potentially making them suitable for pharmaceutical applications where low toxicity is essential [49,50].

**Figure 12:**
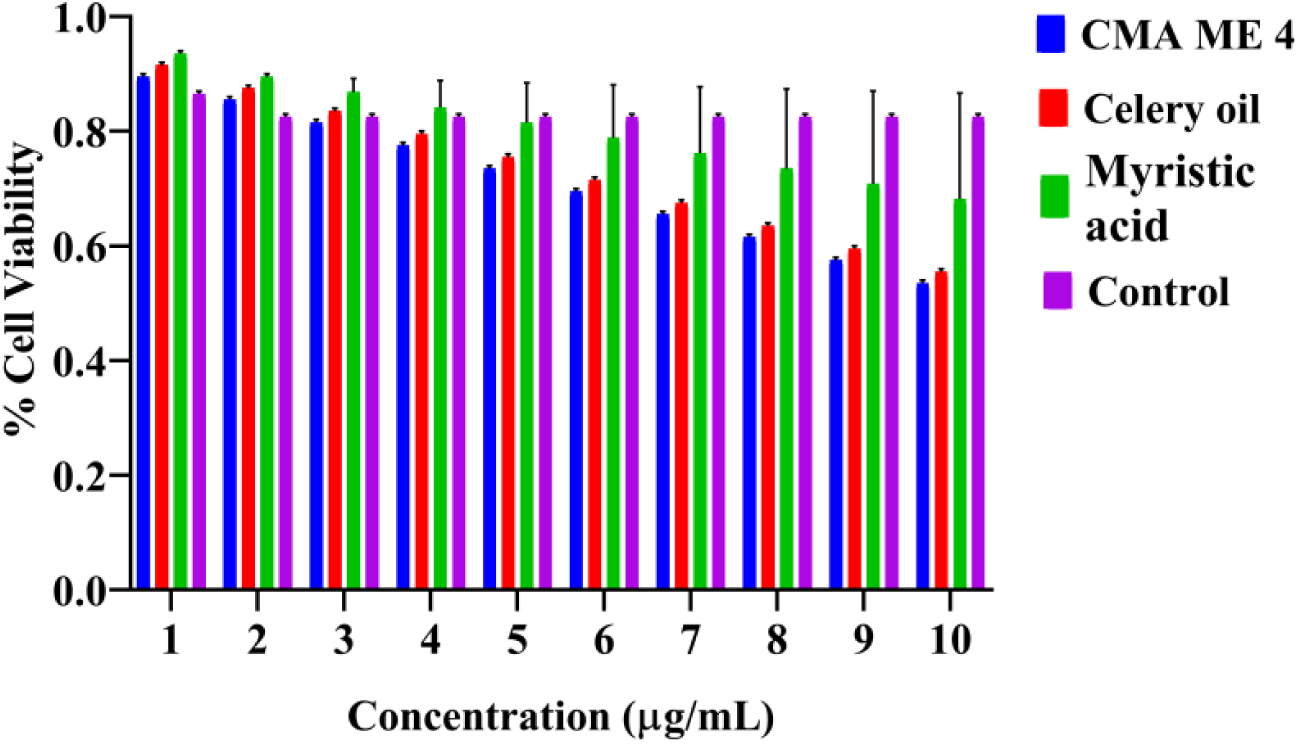
Toxicity analysis using HUVECs cell line

## Conclusion

The celery oil and myristic acid microemulsion has many unique properties. Using a microemulsion system as a carrier to delivery drugs is a feasible and good option. The current work focuses on formulating an effective microemulsion using celery oil and myristic acid, a combined drug preparation. The celery oil which is naturally a good source for numerous health benefits and myristic acid on other hand has good effects in disease. Combining them into a nanoscale carrier to explore its anti oxidant, anti inflammatory, anti diabetic and anti obesity was achieved. The current study successfully tested its anti cancer activity in HeLa and MCF7, for toxicity in HUVECs. Results obtained were efficient in a way for future development of combined therapies with natural and synthetic drugs for targeted delivery for a better future scope.

## Author contribution

**Balaji Govindaswamy –** Conceptualisation, Original draft writing, Data analysis interpretation, Review and editing, Investigation, **Bharathi Ravikrishnan –** Conceptualisation, Data analysis, Review and editing, Supervision, **Dipti Sree panda –** Methodology, Data analysis, Draft writing, **Sarath Perumal –** Data analysis, Methodology, Review and editing

## Acknowledgements

We would like to thank the Guru Nanak College for providing us the instrument, chemicals and space to conduct the research. We also like to thank Indian Institute of Technology Madras and Madurai Kamaraj University for providing necessary instrument for analysis.

